# A Multiple Peptides Vaccine against nCOVID-19 Designed from the Nucleocapsid phosphoprotein (N) and Spike Glycoprotein (S) via the Immunoinformatics Approach

**DOI:** 10.1101/2020.05.20.106351

**Authors:** Sahar Obi Abd Albagi, Mosab Yahya Al-Nour, Mustafa Elhag, Asaad Tageldein Idris Abdelihalim, Esraa Musa Haroun, Mohammed Elmujtba Adam Essa, Mustafa Abubaker, Mohammed A. Hassan

**Author notes:** **Corresponding Author:** Sahar Obi Abd Albagi E.mai.

## Abstract

Due to the current COVID-19 pandemic, the rapid discovery of a safe and effective vaccine is an essential issue, consequently, this study aims to predict potential COVID-19 peptide-based vaccine utilizing the Nucleocapsid phosphoprotein (N) and Spike Glycoprotein (S) via the Immunoinformatics approach. To achieve this goal, several Immune Epitope Database (IEDB) tools, molecular docking, and safety prediction servers were used. According to the results, The Spike peptide peptides **SQCVNLTTRTQLPPAYTNSFTRGVY** is predicted to have the highest binding affinity to the B-Cells. The Spike peptide **FTISVTTEI** has the highest binding affinity to the MHC I HLA-B1503 allele. The Nucleocapsid peptides **KTFPPTEPK** and **RWYFYYLGTGPEAGL** have the highest binding affinity to the MHC I HLA-A0202 allele and the three MHC II alleles HLA-DPA1*01:03/DPB1*02:01, HLA-DQA1*01:02/DQB1- *06:02, HLA-DRB1, respectively. Furthermore, those peptides were predicted as non-toxic and non-allergen. Therefore, the combination of those peptides is predicted to stimulate better immunological responses with respectable safety.

## Introduction

Human coronaviruses (HCoVs) including the Severe acute respiratory syndrome coronavirus 2 (SARS-CoV-2, COVID-19) are enveloped, positive-sense, single-stranded polyadenylated RNA viruses belong to the Coronaviridae family. They cause systemic and respiratory zoonotic diseases [1].

The SARS-CoV-2 is a novel strain detected firstly in the city of Wuhan, the Republic of China in December 2019 [2]. It causes fever, cough, dyspnea, bilateral infiltrates on chest imaging and may progress to Pneumonia [3]. The COVID-19 is characterized by rapid spreading; “As the 27 Feb, it is reported in 47 countries, causing over 82,294 infections with 2,804 deaths” [4] and till the 5th May, more than 3. 5 million positive cases and 0.25 million deaths have been identified globally [5]. Unfortunately, until now COVID-19 has no effective antiviral drug for the treatment or vaccine for the prevention, hence extensive researches should be conducted on the development of safe and effective vaccines and antiviral drugs [4].

To develop a safe and effective COVID-19 vaccine rapidly, the WHO recommended that “we must test all candidate vaccines until they fail to ensure that all of them have the chance of being tested at the initial stage of development”. Ensuing this point, recently, there are over 120 proposed vaccines. Six of them in the clinical evaluation and 70 in pre-clinical evaluation [6]. The vaccine development is achieved by multiple approaches including the Inactivated, Live-attenuated, Non-replicating viral vector, DNA, RNA, Recombinant proteins, and Peptide-based vaccines. “As of 8 April 2020, the global COVID-19 vaccine R&D landscape confirmed 78 active candidate vaccine” [7].

Consistent with global efforts, this study aims to predict potential COVID-19 Peptide-based vaccine utilizing the Nucleocapsid phosphoprotein (N) and Spike Glycoprotein (S) via the Immunoinformatics approach. Due to the respectable antigenicity of the Nucleocapsid and Spike Glycoprotein, they are appropriate targets for vaccine design [8]. The peptide vaccines are sufficient to stimulate cellular and humoral immunity without allergic responses [9]. They are” safe, simply produced, stable, reproducible, cost-effective” [10], and permits a broad spectrum of immunity [9], consequently, they are the targets for this study as well as they utilized in multiple studies concerning COVID-19 vaccine [11-14].

## Materials and Methods

### Protein Sequence Retrieval

A total of 100 COVID-19 Nucleocapsid phosphoprotein (N) and Spike Glycoprotein (S) were retrieved from the National Center for Biotechnology Information (NCBI) database [15] as FASTA format in March 2020. The sequences with their accession number are listed in the Supplementary file S1.

The protein sequences of MHC I alleles HLA-A*02:01, HLA-B15:03, HLA-C*12:03, and MHC II alleles HLA-DPA1*01:03/DPB1*02:01, HLA-DQA1*01:02/DQB1*06:02, HLA- DRB1 were obtained from the Immuno Polymorphism Database (IPD-IMGT/HLA) [16].

### Multiple Sequences Alignment

The retrieved COVID-19 Nucleocapsid phosphoprotein (N) and Spike Glycoprotein (S) sequences were aligned using the ClustalW algorithm [17] on the BioEdit software version 7.2.5 [18] to identify the conserved regions between sequences.

### B-Cells Peptides Prediction

The B-Cells peptides were predicted from the conserved regions using the linear Epitope Prediction tool “BepiPred-test” on the Immune Epitope Database (IEDB) [19] at the default threshold value −0.500. To predict the epitopes accurately, a combination between the hidden (Parker and Levitt) method and the Markov model (HMM) [20] was used.

### The Surface Accessibility Prediction

The Surface Accessibility of B-Cells Peptides was predicted via the Emini Surface Accessibility tool [21] on the IEDB [19] at the default threshold holding value.

### The Antigenic Sites Prediction

To identify the antigenic sites within the Nucleocapsid phosphoprotein and Spike Glycoprotein, the Kolaskar and Tongaonker method’s on the IEDB [19] at the default threshold value.

### T-Cell Peptides Prediction

To predict the interaction with different MHC I alleles, the Major Histocompatibility Complex class I (MHC I) binding prediction tool on the IEDB [19] was used. All peptide length was set as 9amino acid. To predict the binding affinity, the Artificial Neural Network (ANN) prediction method was selected with a half-maximal inhibitory concentration (IC50) value of less than 100.

In contrast, to predict the interaction with different MHC II alleles, The Major Histocompatibility Complex class II (MHC II) binding prediction was used. To predict the binding affinity, the NN align algorithm was selected with an IC50 value of less than 500. The Human allele reference sets (HLA DR, DP, and DQ) were included in the prediction.

### The Population Coverage Prediction

To predict the percentage of peptides binding with various MHC I and MHC II alleles that cover the world population, the population coverage tool on the IEDB [19] was used.

### Allergicity and Toxicity Prediction

To predict the peptides’ allergicity, the AllergenFP v.1.0 [22] and AllerCatPro v. 1.7 [23] servers were used. In contrast, to predict the peptides’ toxicity, the ToxinPred server [24] was used.

### 3D Structure Modeling and Visualization

To model the 3D structure of the Nucleocapsid, Spike, and MHC molecules, the SWISS-MODEL server [25], and the Phyre2 web portal [26] were used. To visualize the modeled structures, the USCF Chimera 1.8 software [27] was used.

### Molecular Docking Study

The predicted peptides were docked with MHC I alleles HLA-A*02:01, HLA-B15:03, HLA-C*12:03, and MHC II alleles HLA-DPA1*01:03/DPB1*02:01, HLA- DQA1*01:02/DQB1*06:02, HLA-DRB1. The modeled MHC structures were prepared for the docking via Cresset Flare software [28] at the normal type calculation method. To dock the predicted peptides, the MDockPeP [29] and HPEPDOCK [30-34] servers were used. The predicted peptides were submitted as amino acid sequences. The 2D and 3D interactions were visualized using the PoseView [35] at the ProteinPlus web portal [36] and Cresset Flare viewer [28], respectively.

## Results

According to the IEDB [19] prediction, the average binding score for the Nucleocapsid phosphoprotein and Spike Glycoprotein were 0.558, 0.470, respectively. All values equal to or greater than the default threshold were predicted as potential B-cell binders.

The Emini surface accessibility tool predicts the average binding score of the Nucleocapsid phosphoprotein and Spike Glycoprotein as 1.00. All values equal to or greater than the default threshold were predicted to have good surface accessibility. The Kolaskar and Tongaonkar antigenicity prediction tool predicts the average threshold value for the Nucleocapsid phosphoprotein and Spike Glycoprotein as 0.988 and 1.04, respectively. Peptides with values equal to or greater than the average score are considered as antigenic peptides. The minimum and maximum values are listed in Table 1. The Nucleocapsid peptide (**DAYKTFPPTEPKKDKKKKADETQALPQRQKKQQTVTLLPAADLDD**) had the highest antigenicity, surface accessibility, hydrophilicity, and flexibility. In contrast, the Spike peptides (**LGKY**) and (**SQCVNLTTRTQLPPAYTNSFTRGVY**) had the highest scores (Table 2). The location of those peptides within the 3D structure is shown in Figures 1 and 2.

**Table 1.** the average, minimum, and maximum Bepiprd epitopes, surface accessibility, Antigenicity, hydrophilicity, and Flexibility, prediction values.

| Test | Protein | Average Score | Minimum Score | Maximum Score |
| --- | --- | --- | --- | --- |
| **Bepiprd linear epitope prediction** | Nucleocapsid | 0.558 | 0.297 | 0.764 |
|  | Spike Glycoprotein | 0.470 | 0.183 | 0.695 |
| **Emini surface accessibility prediction** | Nucleocapsid | 1.000 | 0.050 | 7.006 |
|  | Spike Glycoprotein | 1.000 | 0.042 | 6.051 |
| **Kolaskar & Tongaonkar Antigenicity** | Nucleocapsid | 0.988 | 0.874 | 1.197 |
|  | Spike Glycoprotein | 1.041 | 0.866 | 1.261 |
| **Parker Hydrophilicity Prediction** | Nucleocapsid | 2.800 | -5.971 | 6.871 |
|  | Spike Glycoprotein | 1.238 | 7.629 | 7.743 |
| **Karplus & Schulz**<br>**Flexibility**<br>**Prediction** | Nucleocapsid | 1.035 | 0.885 | 1.161 |

**Table 2.**
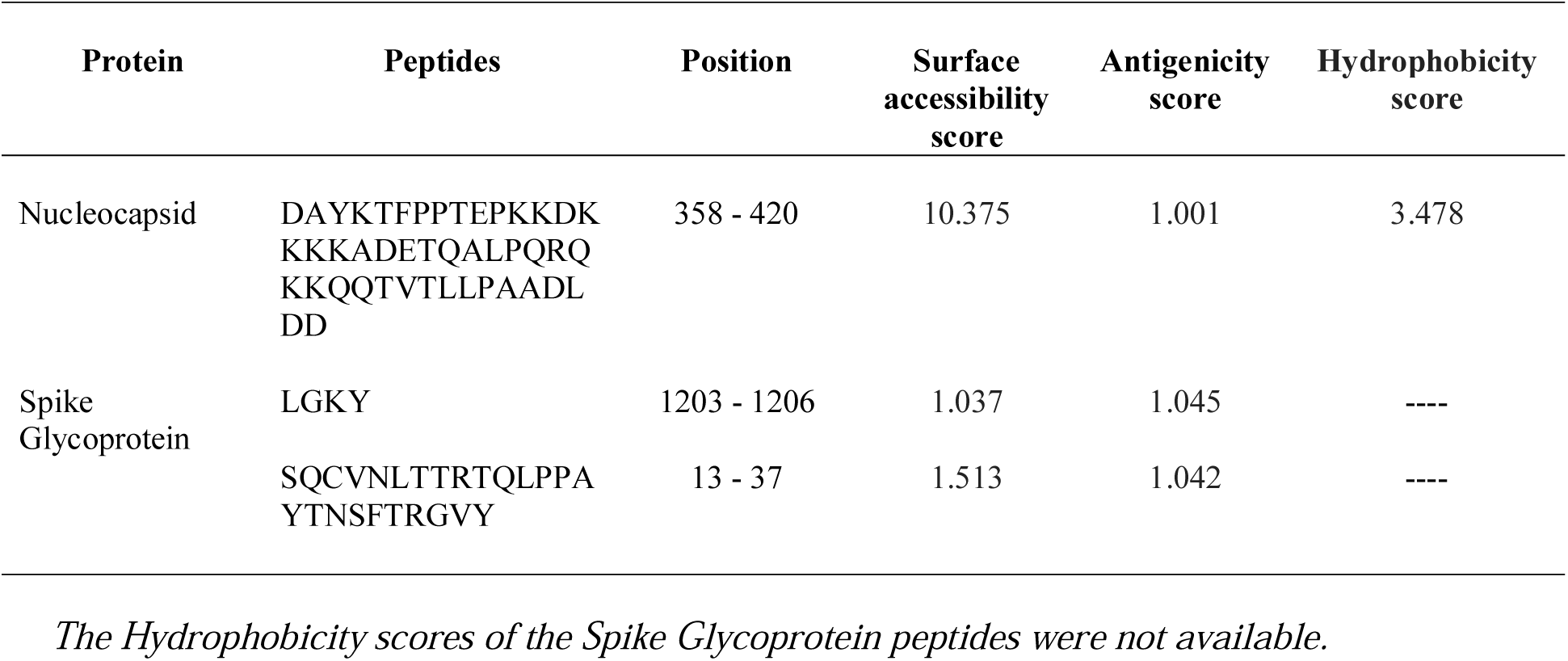
Result of the predicted B cell epitopes.

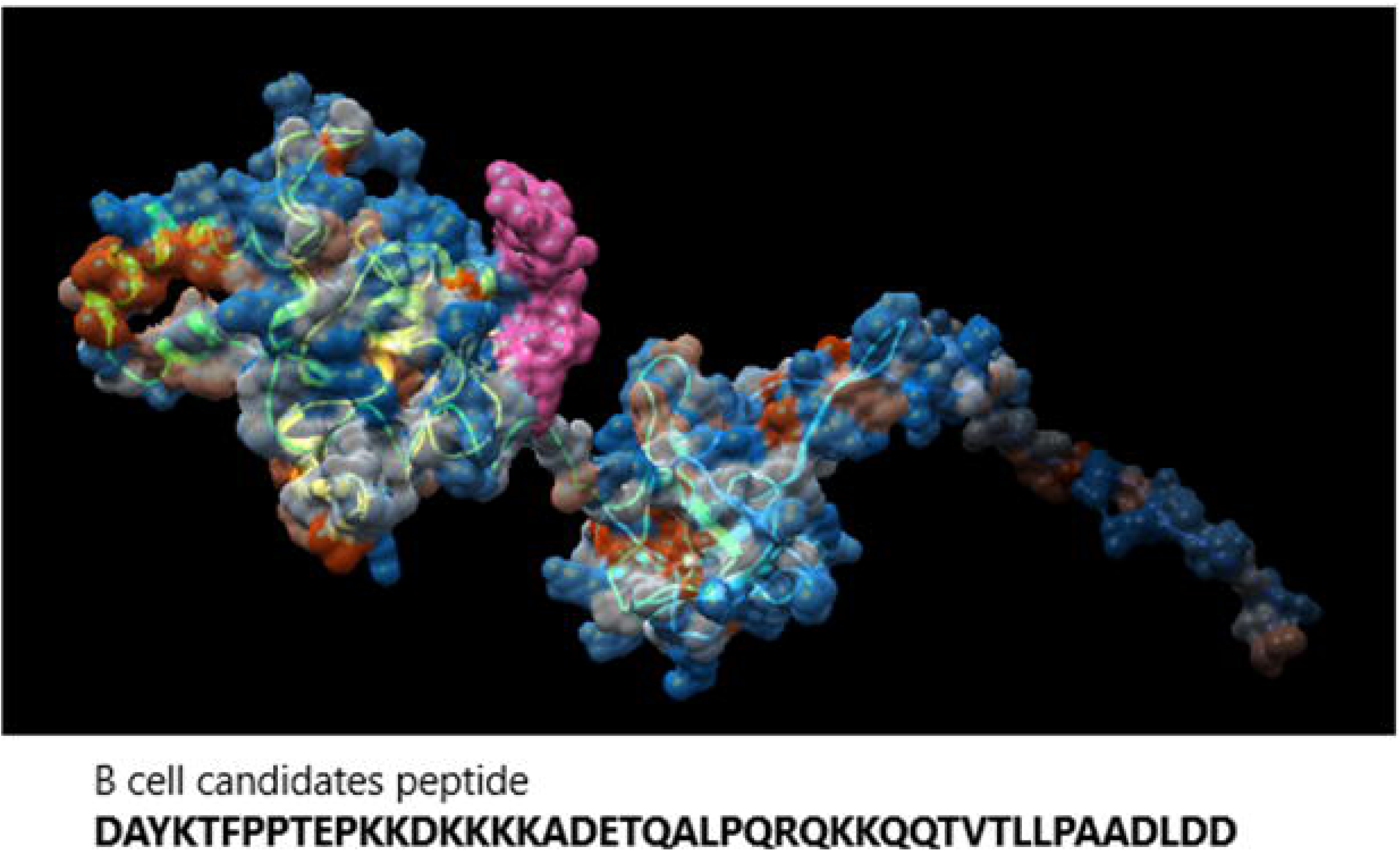

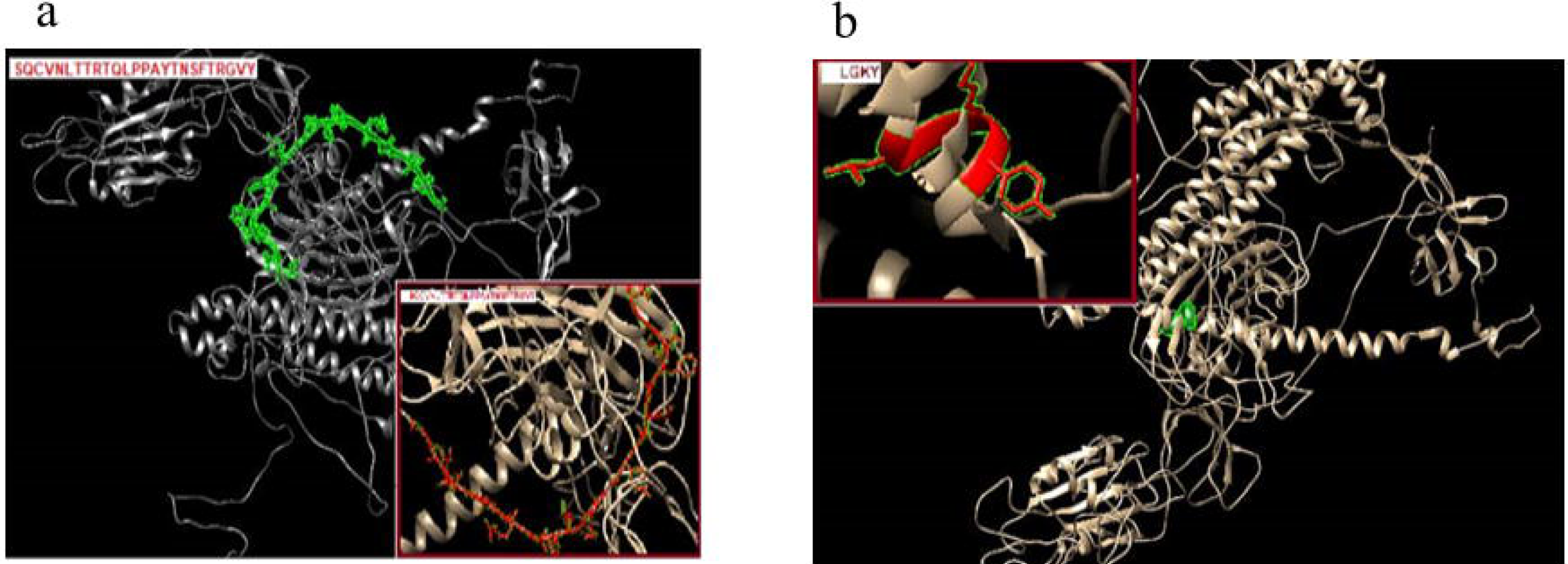

Regarding the cytotoxic T-lymphocyte peptides, the MHC I binding prediction tool predicts 46 peptides from the Nucleocapsid and 192 peptides from the Spike Glycoprotein could interact with the different MHC I alleles. The most promising peptides were listed in Table 3. In contrast, the MHC II binding prediction tool predicts 774 peptides from the Nucleocapsid and 1111 peptides from the Spike Glycoprotein could interact with the different MHC II alleles. The most promising peptides were listed in Table 4. The total population coverage percentage of the promising peptides was above 98%. The remaining MHC I and MHC II peptides are listed in the supplementary file S2. The top predicted MHC I peptides are **KTFPPTEPK** from the Nucleocapsid and **FTISVTTEI, MIAQYTSAL** from the Spike Glycoprotein. The top predicted MHC II peptides are **AALALLLLDRLNQLE, ALALLLLDRLNQLES, PRWYFYYLGTGPEAG, RWYFYYLGTGPEAGL** from the Nucleocapsid and **EVFNATRFASVYAWN, VFRSSVLHSTQDLFL** from the Spike Glycoprotein. The location of representative peptides from them within the 3D structure is shown in Figures 3 and 4.

**Table 3.**
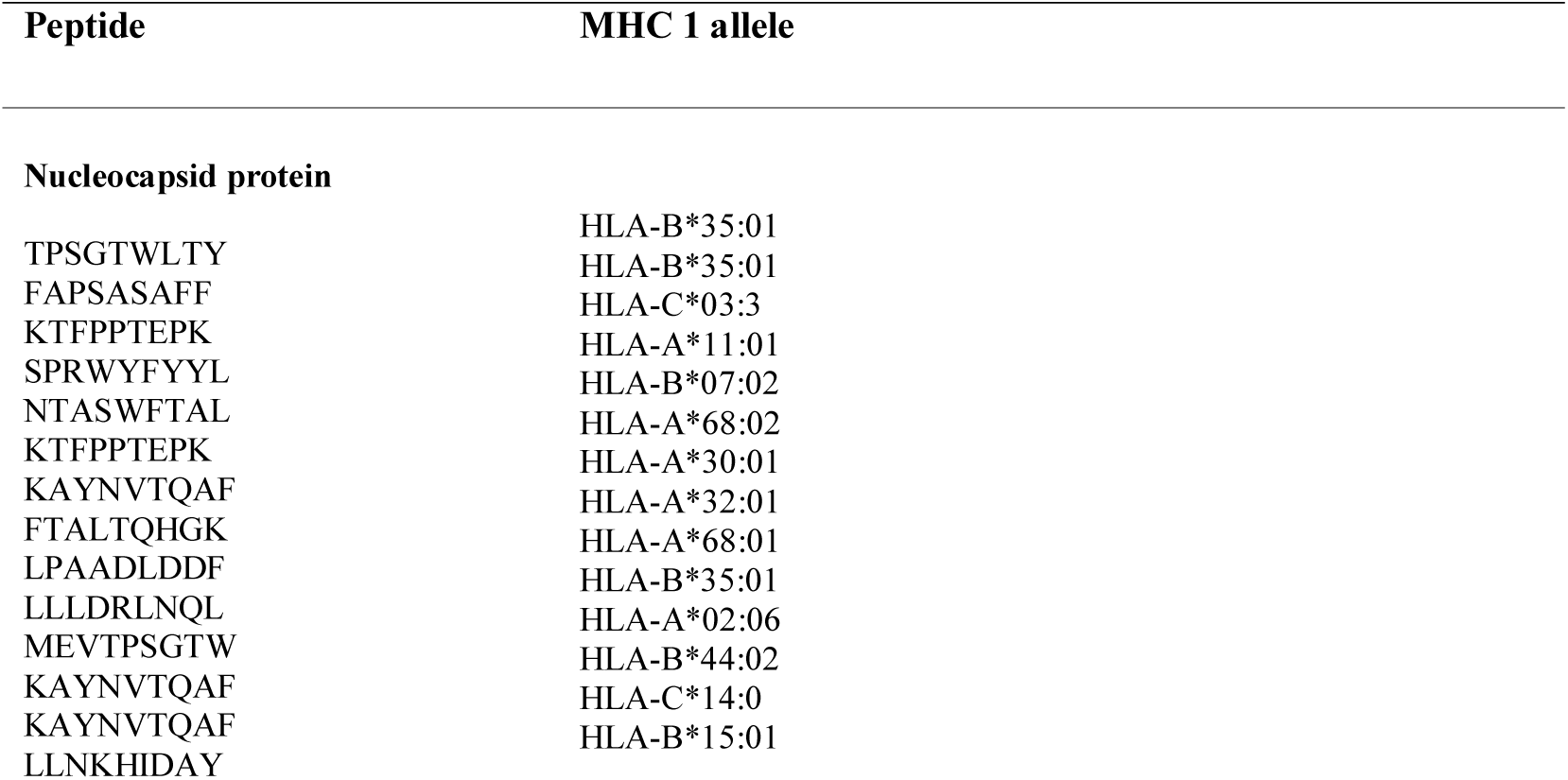
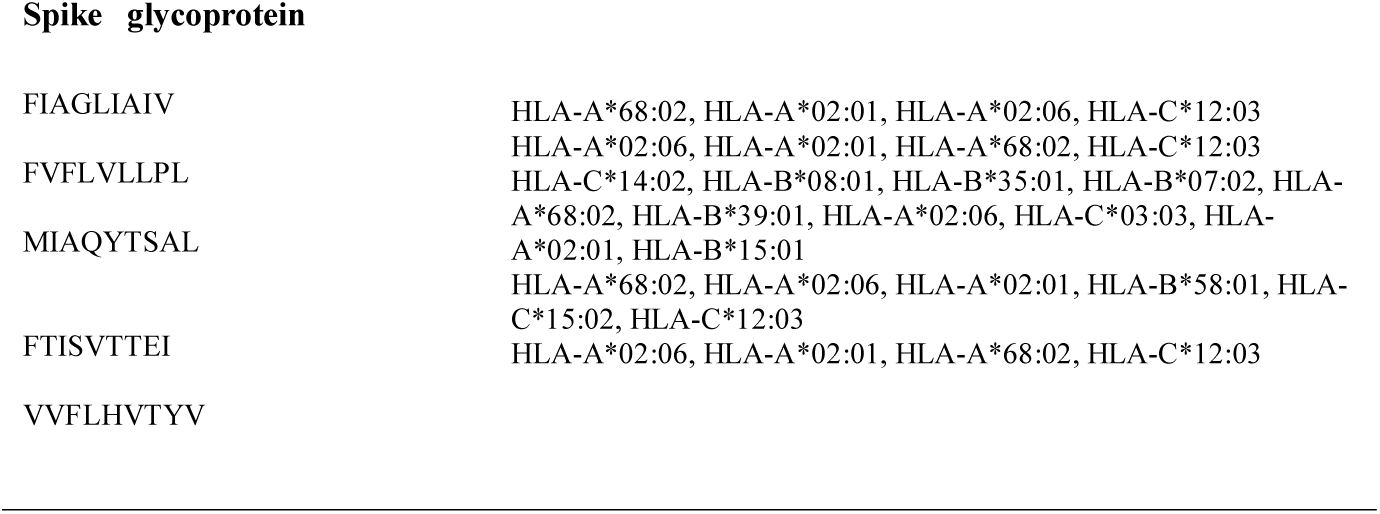
the most promising MHC I peptides and their total population coverage percentage.

**Table 4.**
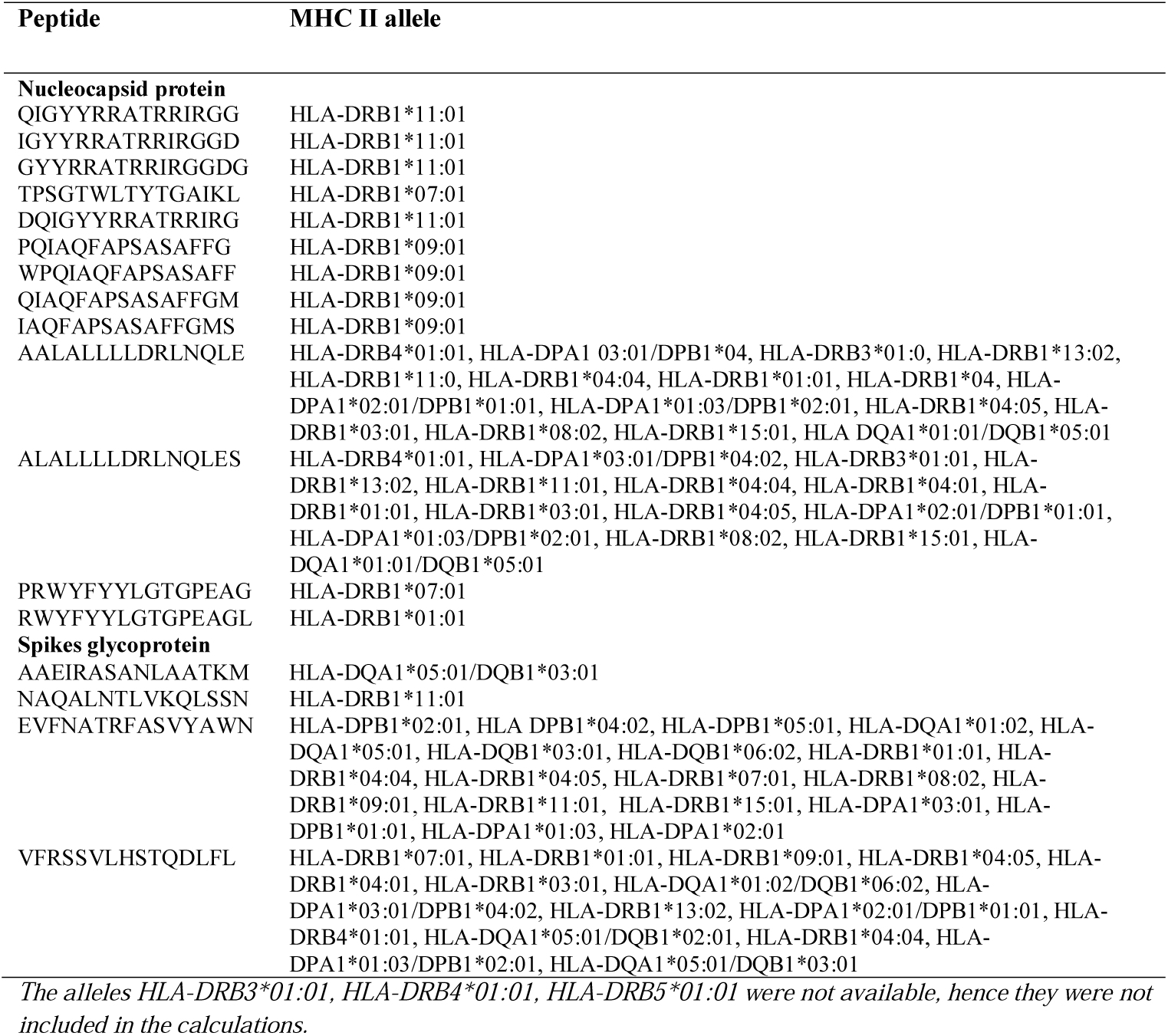
the most promising MHC II peptides and their total population coverage percentage.

**Table 5.** the total population coverage percentage in MHC molecule of the Spike Glycoprotein (S) and Nucleocapsid phosphoprotein (N) peptides

| MHC molecule | Total Population Coverage |
| --- | --- |
| MHC I (Spike) | 99.73% |
| MHC II (Spike) | 99.98% |
| MHC I & II combined (Spike) | 100.0% |
| MHC I (Ncleocapsid) | 98.2% |
| MHC II (Ncleocapsid) | 99.97% |
| MHC I & II combined (Ncleocapsid) | 99.97% |

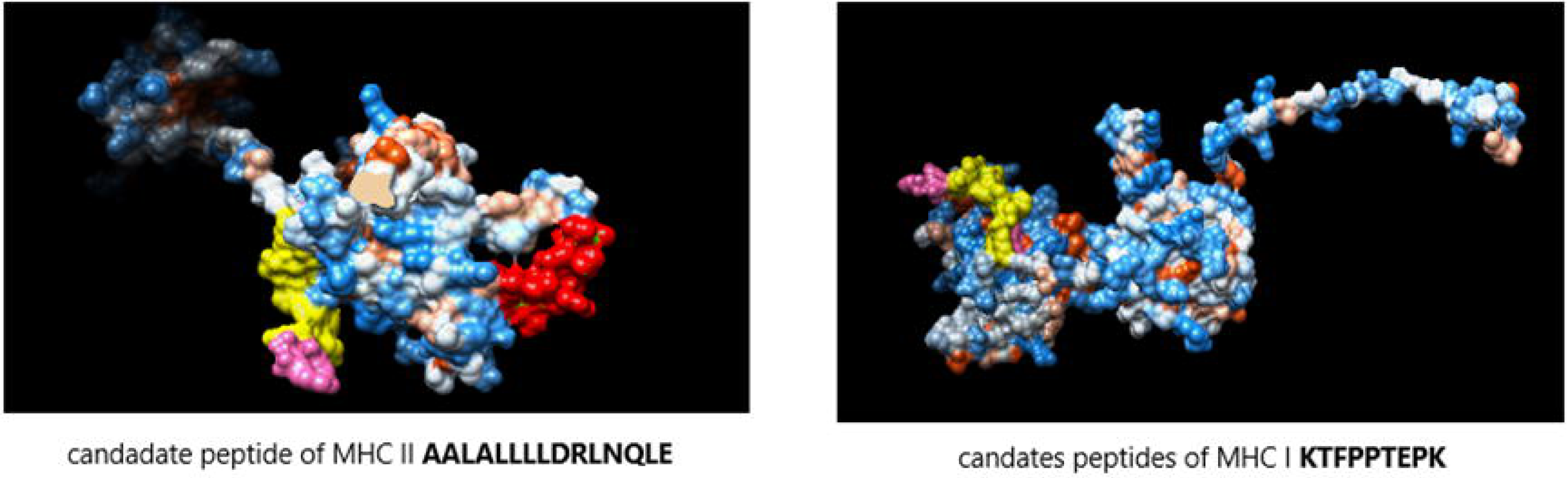

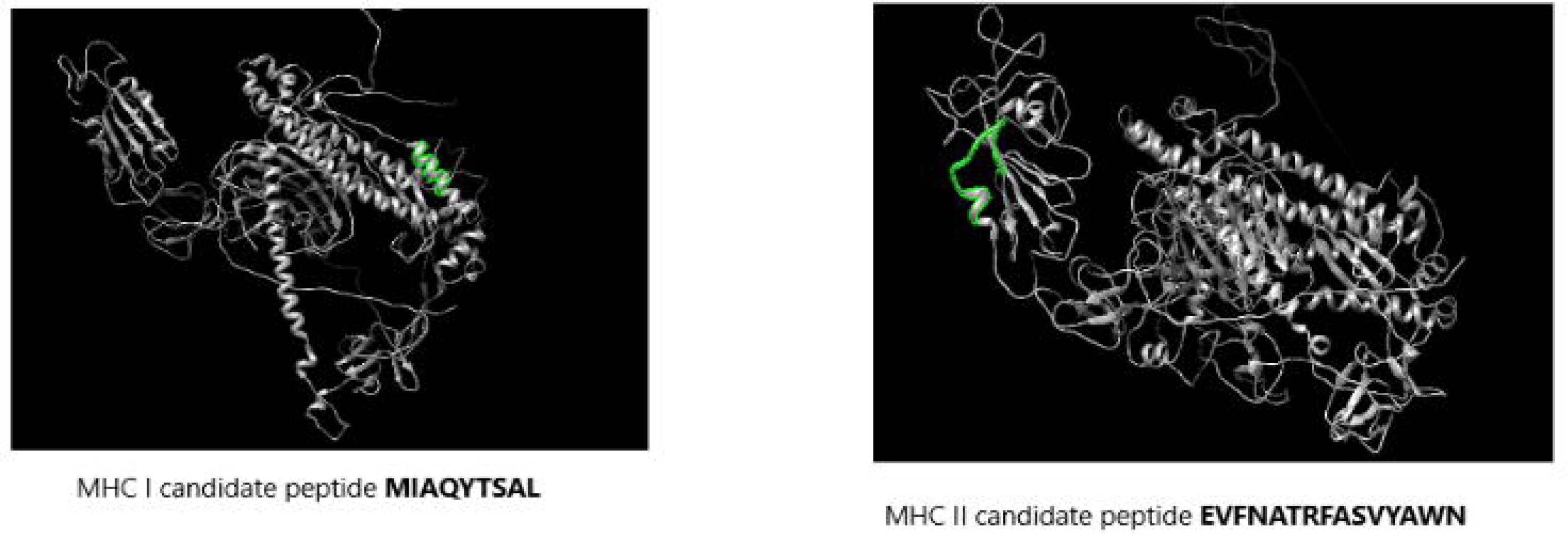

According to the MDockPeP [29] and HPEPDOCK [30] servers prediction, the spike peptide (**FTISVTTEI)** has the highest binding affinity to the MHC I HLA-B1503 allele and the spike peptide (**MIAQYTSAL**) has the highest binding affinity to the MHC I HLA-C1203 allele. The Nucleocapsid peptide (**KTFPPTEPK)** has the highest binding affinity to the MHC I HLA-A0202 allele (Table 6). The 2D and 3D interactions are illustrated in Figures 5 and 6.

**Table 6.** Docking scores of the predicted COVID-19 Spike Glycoprotein (S) and Nucleocapsid phosphoprotein (N) peptides with MHC I alleles HLA-A0202, HLA-B1503, and HLA-C1203.

| Peptide | HLA-A0202 |  | HLA-B1503 |  | HLA-C1203 |  |
| --- | --- | --- | --- | --- | --- | --- |
|  | MDockPeP | HPEPDOCK | MDockPeP | HPEPDOCK | MDockPeP | HPEPDOCK |
| **FTISVTTEI** | -148.1 | -250.625 | **-153.9** | **-229.356** | -134.2 | -215.735 |
| (Spike) |  |  |  |  |  |  |
| **MIAQYTSAL** | -149.3 | -259.707 | -149.4 | -219.145 | **-154.7** | -237.005 |
| (Spike) |  |  |  |  |  |  |
| **KTFPPTEPK** | **-153.9** | -220.876 | -135.8 | -204.661 | -136.3 | -193.690 |
| (Nucleocapsid) |  |  |  |  |  |  |

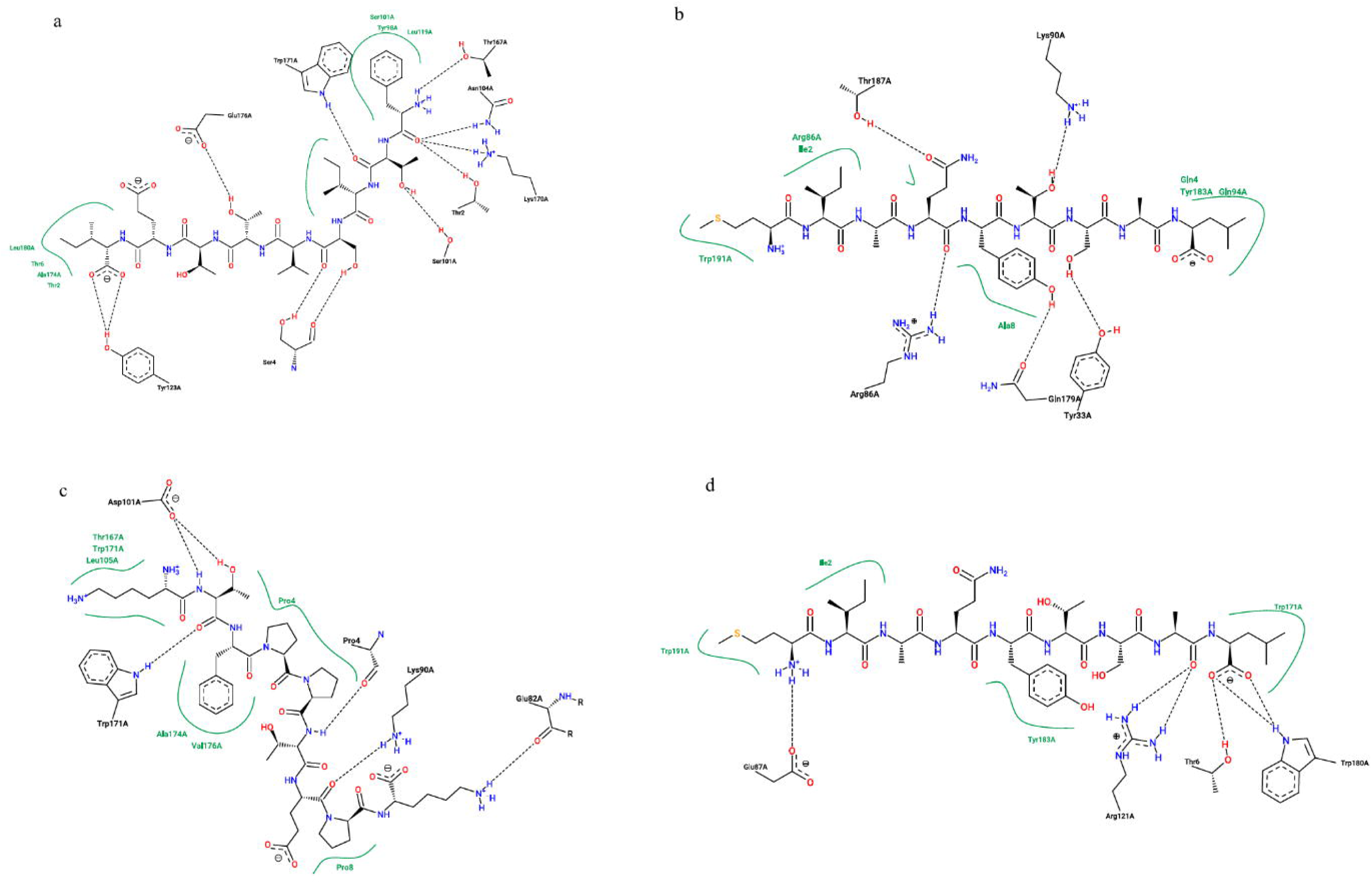

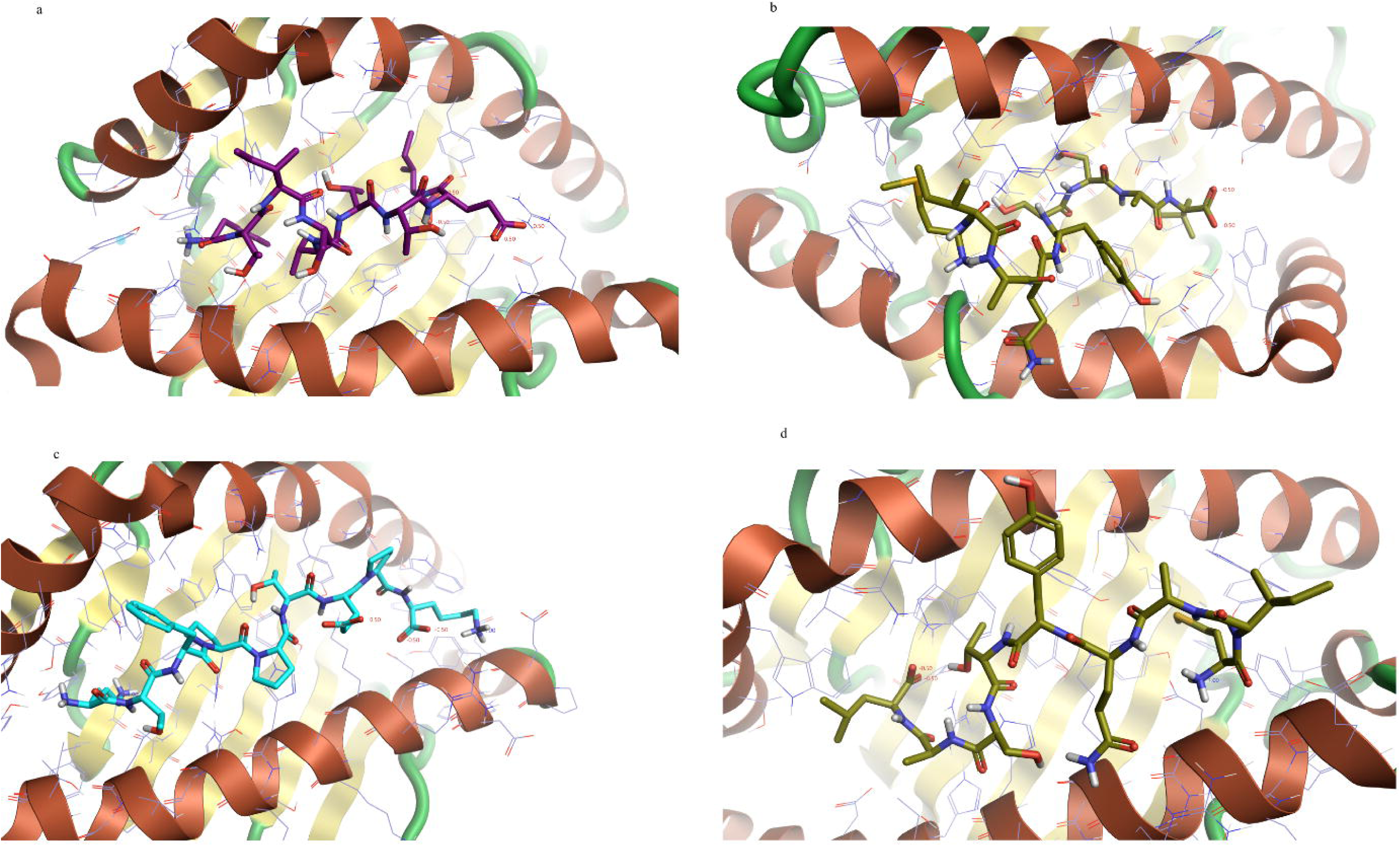

In contrast, the spike peptide (**EVFNATRFASVYAWN**) has the highest binding affinity to the MHC II alleles HLA-DPA1*01:03/DPB1*02:01, HLA-DQA1*01:02/DQB1*06:02, and HLA-DRB1. The Nucleocapsid peptide (**RWYFYYLGTGPEAGL**) has the highest binding affinity to the MHC II allele HLA-DQA1*01:02/DQB1*06:02 and the Nucleocapsid peptides (**RWYFYYLGTGPEAGL**) (**PRWYFYYLGTGPEAG**) and has a higher binding affinity to the MHC II alleles HLA-DPA1*01:03/DPB1*02:01 and HLA-DRB1 (Table 7). The 2D and 3D interactions are shown in Figures 7 and 8.

**Table 7.**
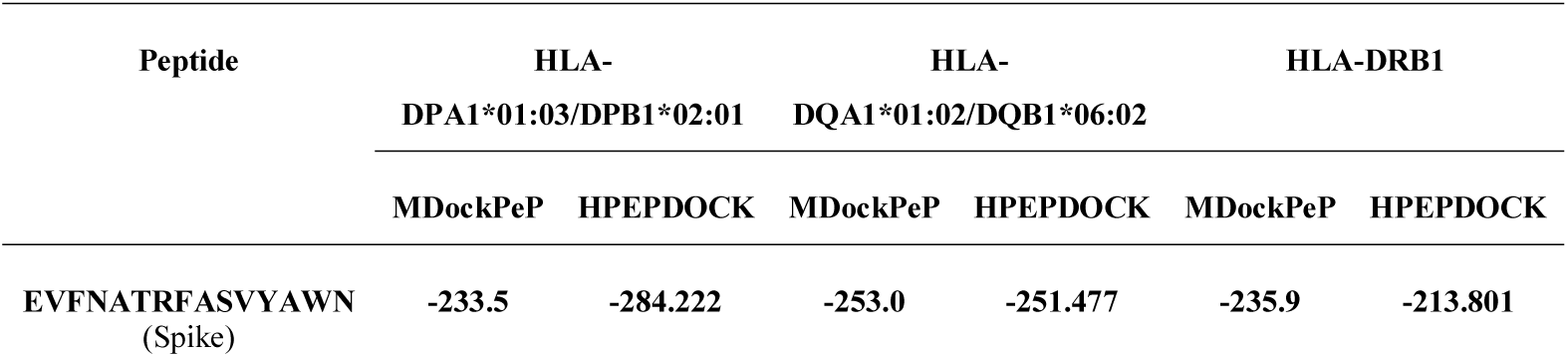
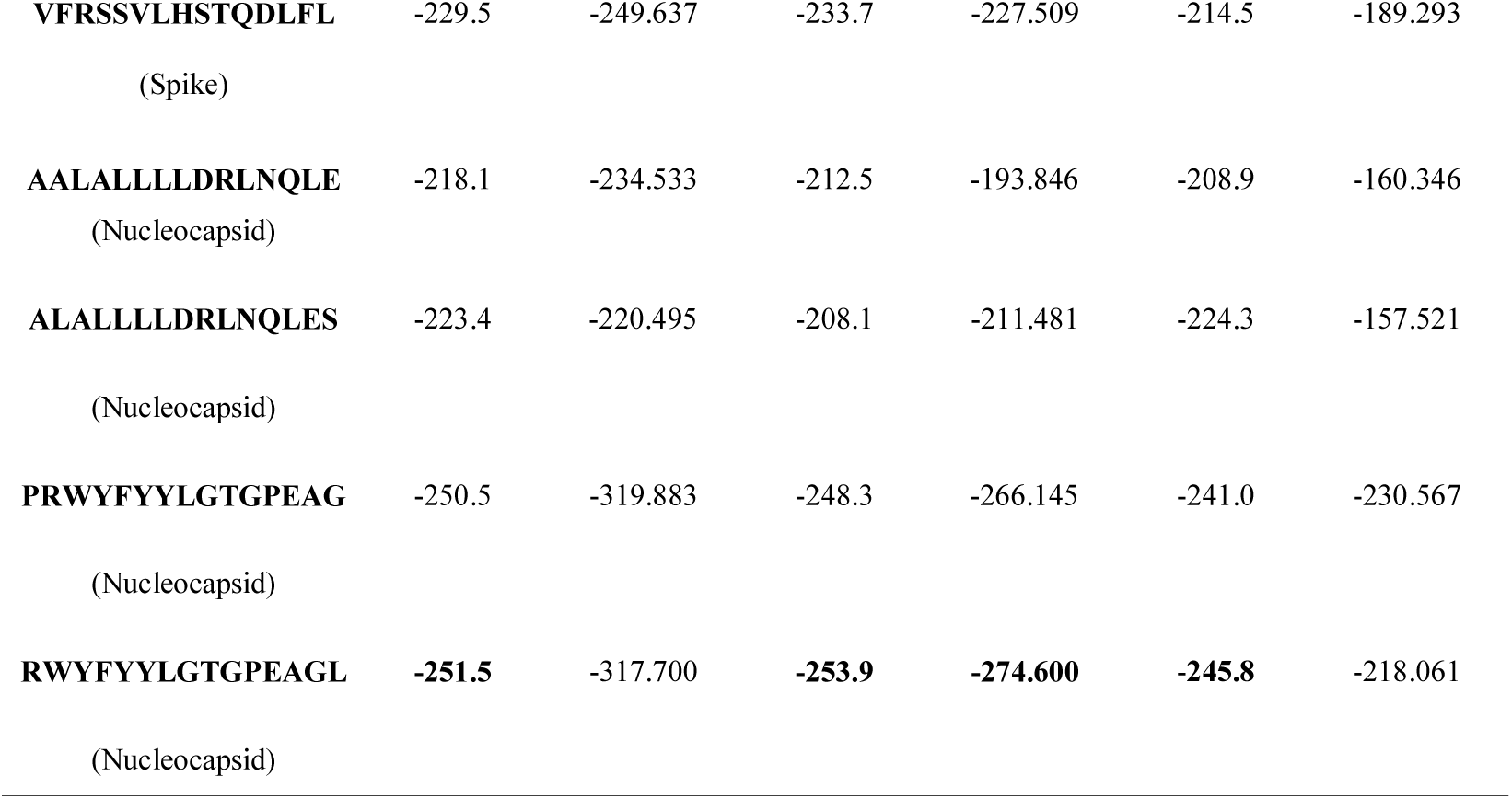
Docking scores of the predicted COVID-19 Spike Glycoprotein (S) and Nucleocapsid phosphoprotein (N) peptides with MHC II alleles HLA-DPA1*01:03/DPB1*02:01, HLA- DQA1*01:02/DQB1*06:02, and HLA-DRB1.

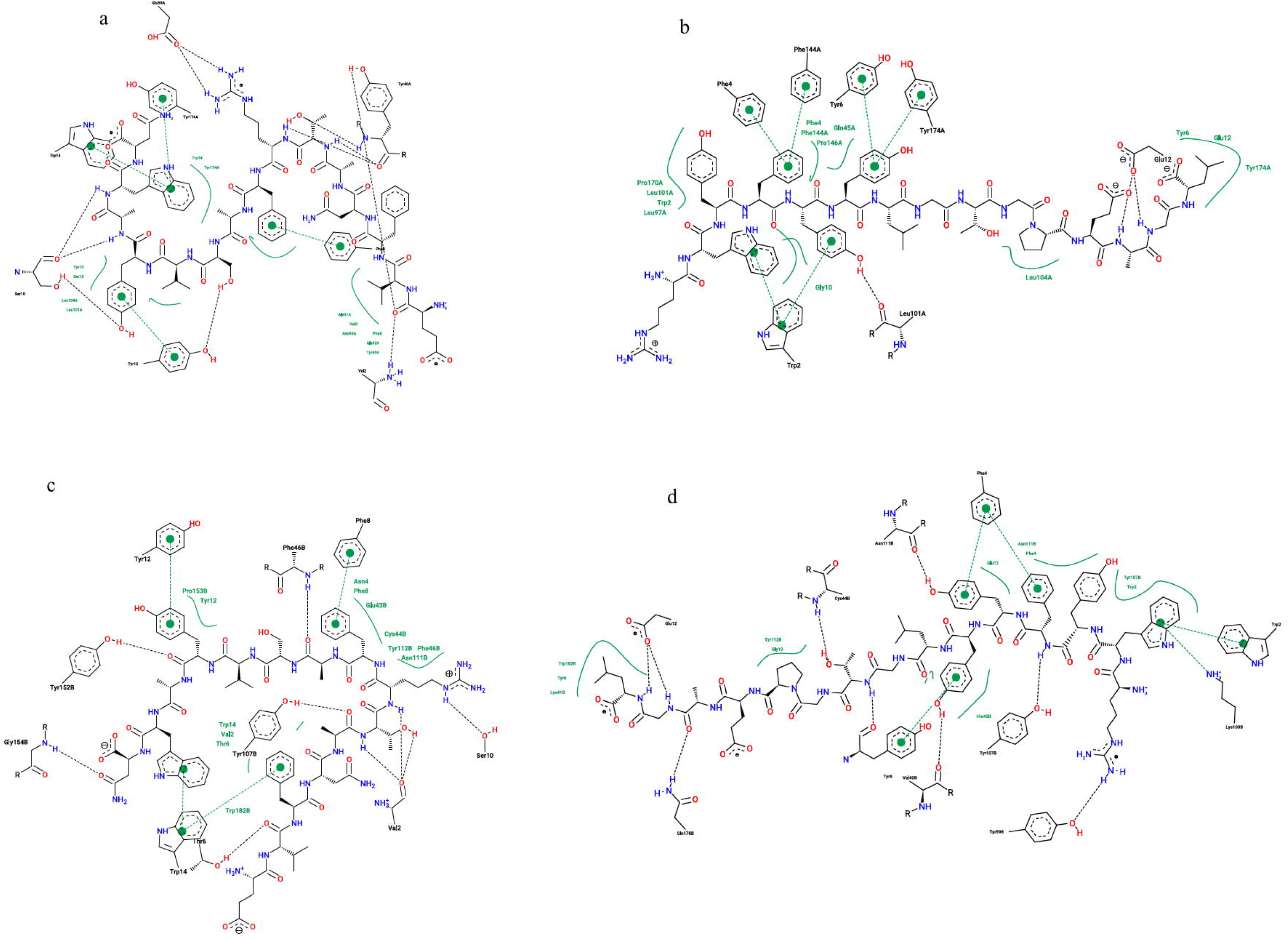

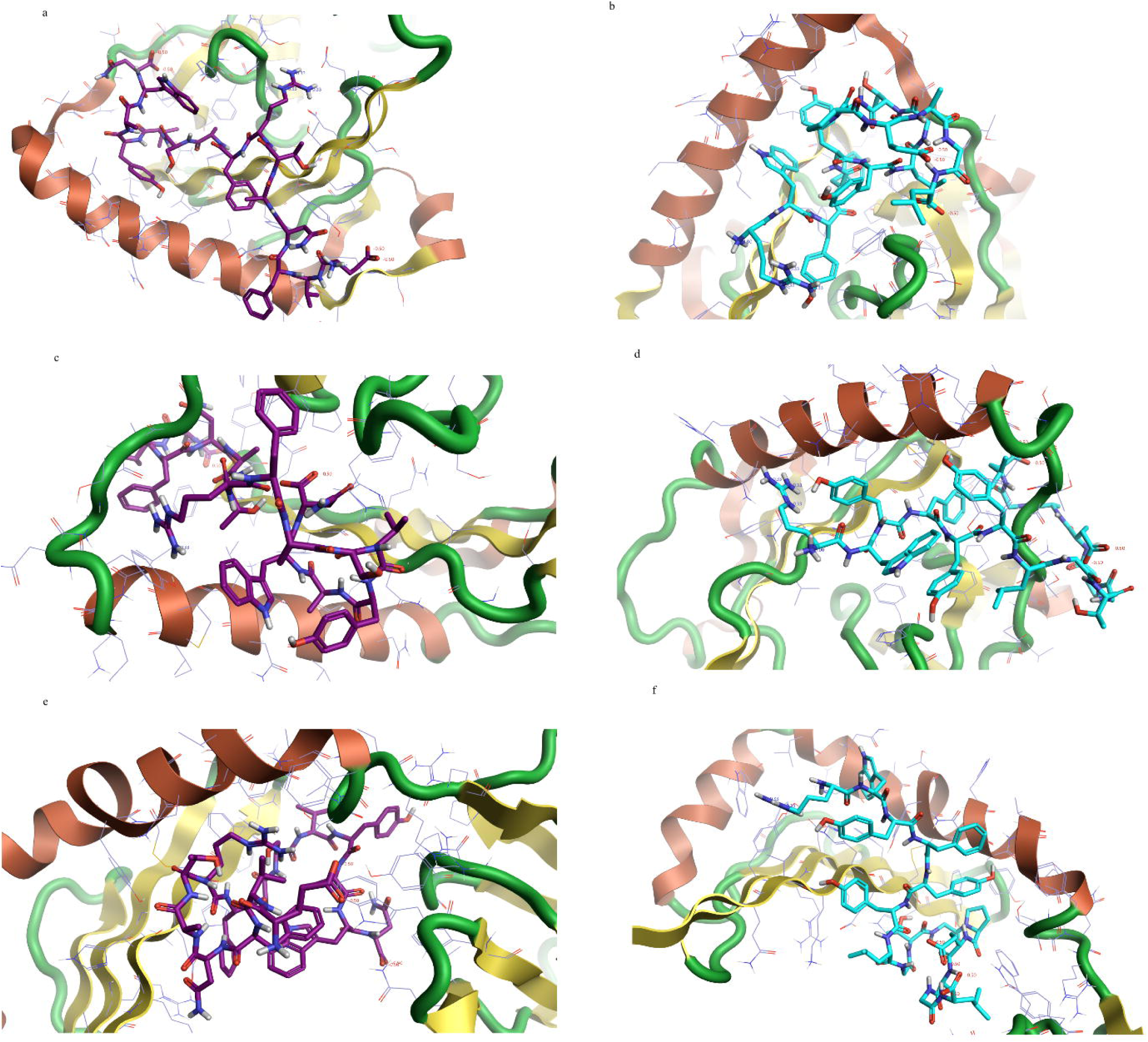

According to the AllergenFP v.1.0 [12], AllerCatPro v. 1.7 [13], and ToxinPred servers [14], all the predicted peptides except the spike peptide (**EVFNATRFASVYAWN**) were Non-allergen and Non-Toxin (Tables 8 and 9).

**Table 8.** Results of MHC I peptides Allergicity and Toxicity prediction.

| Peptide | Allergicity |  | Toxicity |
| --- | --- | --- | --- |
|  | AllergenFP v.1.0 | AllerCatPro v. 1.7 |  |
| **FTISVTTEI**<br>(Spike) | Non-allergen | No evidence | Non-Toxin |
| **MIAQYTSAL**<br>(Spike) | Non-allergen | No evidence | Non-Toxin |
| **KTFPPTEPK**<br>(Nucleocapsid) | Non-allergen | No evidence | Non-Toxin |

**Table 9.**
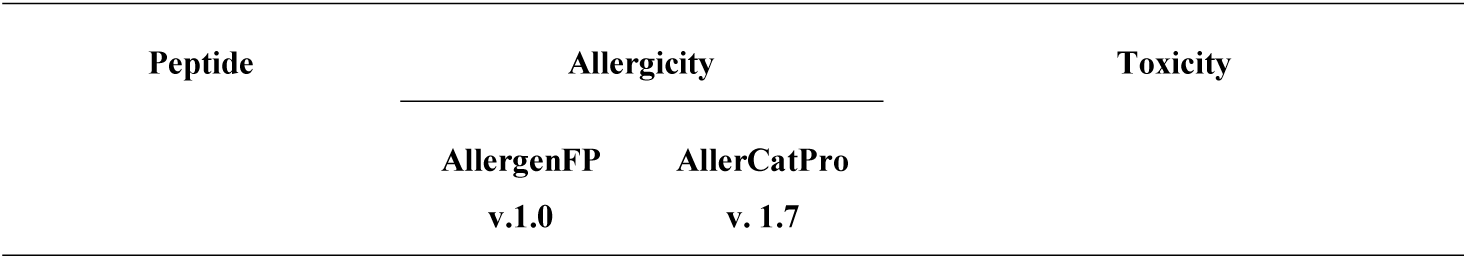
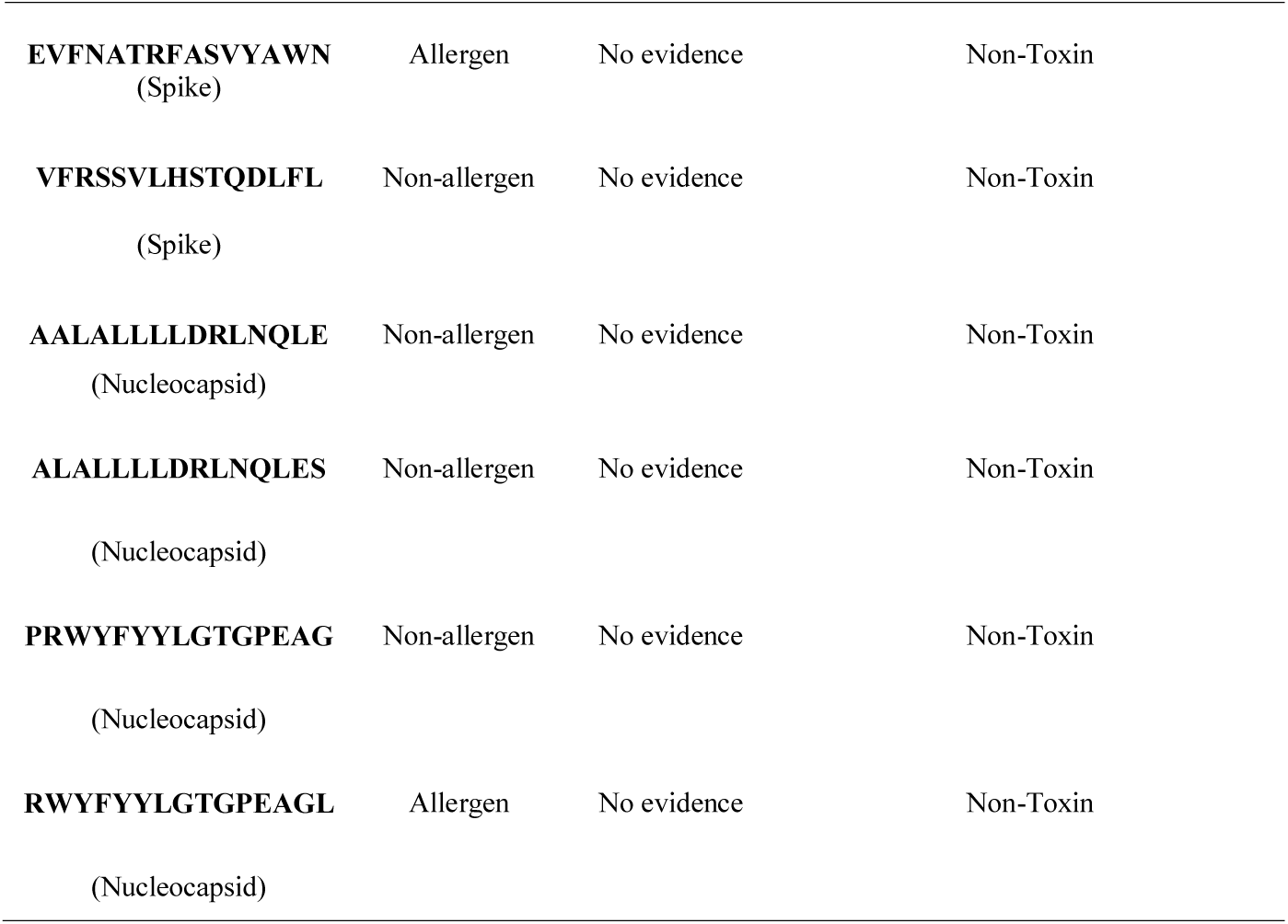
Results of MHC II peptides Allergicity and Toxicity prediction

## Discussion

Due to the current COVID-19 pandemic, the rapid discovery of a safe and effective vaccine is an essential issue [37].

Since the successful vaccine relies on the selection of the most antigenic parts and the best approaches [38], COVID-19 Nucleocapsid phosphoprotein (N) and Spike Glycoprotein (S) were selected to design a peptide vaccine. The antigenicity of the Nucleocapsid and Spike is well predicted [8], and the advantages of the peptide vaccines are well established [9, 10].

The peptide design via the Immunoinformatics approach is achieved through multiple steps including the prediction of; B-Cells and T-cell Peptides, the surface accessibility, antigenic sites, and the Population coverage. After the selection of the candidate peptides, their interaction with the MHC molecules is simulated and their safety is predicted [39].

Regarding the B-Cells peptides prediction, the successful candidates must pass the threshold scores in the Bepipred, Parker hydrophilicity, Kolaskar and Tongaonkar antigenicity, as well as Emini surface accessibility tests [40]. The IEDB Bepipred test [19] on the Nucleocapsid showed that eleven peptides were predicted, however, the peptide **DAYKTFPPTEPKKDK-KKKADETQALPQRQKKQQTVTLLPAADLDD** was the only one that passed all the tests. In contrast, the IEDB Bepipred test [18] on the Spike showed that forty-two peptides were predicted, but the peptides **SQCVNLTTRTQLPPAYTNSFTRGVY** and **LGKY** the only two that passed all the tests. As the length of effective B-cell peptides varies from 5- 30 amino acids [41], the peptide **LGKY** is too short and the peptide **DAYKTFPPTEPKKDK- KKKADETQALPQRQKKQQTVTLLPAADLDD** is too long. Consequently, the Spike peptide peptides **SQCVNLTTRTQLPPAYTNSFTRGVY** is predicted to have the highest binding affinity to the B-Cells (Table 2).

Concerning the T-Cells peptides prediction, the test measures the peptides’ binding affinity to the MHC molecules [19]. The available MHC I alleles HLA A, HLA B, HLA C, HLA E, and MHC II alleles HLA-DR, HLA-DQ, and HLA-DP were used. The MHC I IEDB tests [19] on the Spike glycoprotein predicted 192 peptides could interact with 2-4 MHC I alleles. The top five are **FIAGLIAIV, FTISVTTEI, FVFLVLLPL, MIAQYTSAL**, and **VVFLHVTYV** with population coverage of 99.73%. The tests on the Nucleocapsid protein predicted 46 peptides could interact with 2-4 MHC II alleles. The top three are **KAYNVTQAF, KTFPPTEPK**, and **LLNKHIDAY** with total population coverage of 98.20% (Table 3).

In contrast, the MHC II test on the Spike glycoprotein predicted 1111 peptides could interact with the main MHC II alleles. The top two are **EVFNATRFASVYAWN** and **VFRSSV- LHSTQDLFL** with total population coverage of 99.55%. The tests on the Nucleocapsid protein predicted 177 could interact with the main MHC II alleles. The top six are **AALALLLLDRLNQLE, ALALLLLDRLNQLES, ALLLLDRLNQLESKM, LALLLL- DRLNQLESK, PRWYFYYLGTGPEAG**, and **RWYFYYLGTGPEAGL** with total population coverage up to 98.6% (Table 4).

The results of collective The IEDB tests [19] revealed that the Spike glycoprotein peptides **FTISVTTEI, MIAQYTSAL**, and the Nucleocapsid peptide **KTFPPTEPK** are the most promised MHC1 peptides. On the other hand, the Spike peptides **EVFNATRFASVYAWN, VFRSSVLHSTQDLFL**, and the Nucleocapsid peptides **AALALLLLDRLNQLE, ALALL-LLDRLNQLES, PRWYFYYLGTGPEAG, RWYFYYLGTGPEAGL** are the most promised MHC II peptides.

To stimulate better immunological responses by the predicted peptides, they must interact and bind effectively with the MHC1 and MHC II molecules [42], therefore we must study their interaction with the MHC molecules.

The simulation and prediction of the interaction between the predicted peptides and the MHC molecules are conducted using molecular docking studies that rely on the calculation of the binding free energy. The lowest binding energy scores of the MHC-Peptide complex will indicate the best interaction and the highest stability [43].

To validate the results of molecular docking, MDockPeP [29] and HPEPDOCK [30-34] servers were used. The MDockPeP server predicts the MHC-Peptides interaction by “docking the peptides onto the whole surface of protein independently and flexibly using a novel the conformation restriction in its novel iterative approach. It ranks the docked Peptides via the ITScorePeP scoring function that uses the known protein-peptide complex structures in the calculations” [29]. In contrast, HPEPDOCK uses “a hierarchical flexible peptide docking approach” to predict the MHC-Peptides interaction [30]. The MHC I, HLA-A0202, HLA- B1503, HLA-C1203 was predicted to present the highly conserved SARS-CoV-2 peptides more effectively [44], hence, they were used in molecular docking study.

The molecular docking results showed that the Spike peptide **FTISVTTEI** has the lowest docking energy score with the MHC I HLA-B1503 allele, hence it is predicted to have the highest binding affinity. The Spike peptide **MIAQYTSAL** showed the lowest docking energy score with the MHC I HLA-C1203, consequently, it is predicted to have the highest binding affinity to the MHC I HLA-C1203 allele. In contrast, regarding the MHC I HLA- A0202; the results of the MDockPeP [29] server showed that the Nucleocapsid peptide **KTFPPTEPK** has the lowest docking energy score, but the results of HPEPDOCK [30] server showed the Spike peptide **MIAQYTSAL** has the lowest docking energy score (Table 6).

To illustrate the MHC-Peptide interaction, the PoseView [35] at the ProteinPlus web portal [36] that illustrate the 2D interactions and Cresset Flare viewer [8] that illustrate the 3D interaction were used.

The Spike peptide **FTISVTTEI** interacts with the MHC I HLA-B1503 allele by forming hydrogen bonds with the amino acids Thr2, Ser4, Ser101A, Asn104A, Tyr123A, Thr167A, Lys170A, Trp171A, Glu176A and hydrophobic bonds with the amino acids Thr2, Thr6, Trp98A, Ser101A, Trp171A, Ala174A, Leu180A.

The Spike peptide **MIAQYTSAL** interacts with the MHC I HLA-C1203 allele by forming hydrogen bonds with the amino acids Tyr33A, Arg86A, Lys90A, Gln179A, Thr187A and hydrophobic bonds with the amino acids Ile2, Gln4, Gln94A, Ala8, Arg86A, Tyr183A, Trp191A. In comparison, it interacts with the MHC I HLA-A0202 allele by forming hydrogen bonds with the amino acids Thr6, Glu87A, Arg121A, Trp180A, and hydrophobic bonds with the amino acids Ile2, Trp171A, Tyr183A, Trp191A. It forms six hydrogen bonds and seven hydrophobic bonds with the MHC I HLA-C1203 allele that is more than its bonds with the MHC I HLA-A0202 allele. This finding indicates the higher binding affinity of Spike peptide **MIAQYTSAL** to the MHC I HLA-C1203 allele and supports the MdockPeP [3] server score (Table 6 and Figures 5, 6).

The Nucleocapsid peptide **KTFPPTEPK** interacts with the MHC I HLA-A0202 allele by forming hydrogen bonds with the amino acids Pro4, Glu82A, Lys90A, Asp101A, Trp171A and hydrophobic bonds with the amino acids Pro4, Pro8, Leu105A, Thr167A, Trp171A, Ala174A, Val176A (Figures 5 and 6).

Among the reported MHC I alleles, the HLA-B1503 allele was predicted to have “the greatest ability to present the highly conserved SARS-CoV-2 peptides” [44], therefore, the Spike peptide **FTISVTTEI** is predicted to make the highest response, since the binding with the MHC I stimulates the natural killer and the cytotoxic T cells [45].

Regarding the interaction with the MHC II molecule, the Spike peptide **EVFNATRFASVYAWN** showed the lowest docking energy score with the three MHC II alleles HLA-DPA1*01:03/DPB1*02:01, HLA-DQA1*01:02/DQB1*06:02, and HLA-DRB1, hence it is predicted to have the highest binding affinity to the three alleles. Hence, it predicted to stimulate the CD4+ (helper) T cells more effectively, since the MHC II molecule presents the antigenic peptides to the CD4+ (helper) T cells [46].

On the contrary, the Nucleocapsid peptide (**RWYFYYLGTGPEAGL**) showed lower docking energy scores with the MHC II allele HLA-DQA1*01:02/DQB1*06:02. The results of the MDockPeP [29] server showed that the peptide (**PRWYFYYLGTGPEAG**) has the lowest docking energy score, however, the results of HPEPDOCK [30] server differed from it in the MHC II alleles HLA-DPA1*01:03/DPB1*02:01 and HLA-DRB1 (Table 7).

The Spike peptide (**EVFNATRFASVYAWN**) interacts with HLA- DPA1*01:03/DPB1*02:01 allele by forming hydrogen bonds with the amino acids Val2, Ser10, Tyr12, Tyr40A, Glu59A, hydrophobic bonds with the amino acids Val2, Phe8, Tyr40A, Ala41A, Ala42A, Asn93A, Leu101A, Leu104A, Trp14, Tyr174A, and pi-pi bonds with the aromatic amino acid Phe8, Ser10, Tyr12, Trp14, Tyr174A. It interacts with the HLA-DRB1 allele by forming hydrogen bonds with the amino acids Val2, Thr6, Ser10, Phe46B, Tyr107B, Tyr152B, Gly154B, hydrophobic bonds with the amino acids Val2, Asn4, Thr6, Phe8, Tyr12, Trp14, Glu43B, Cys44B, Phe46B, Asn111B, Tyr112B, Pro153B, Trp182B, and pi-pi bonds with the aromatic amino acid Phe8, Tyr12, Trp14. Its interaction with the HLA-DQA1*01:02/DQB1*06:02 allele was not obtained from the server.

The Nucleocapsid peptide (**RWYFYYLGTGPEAGL**) interacts with HLA- DPA1*01:03/DPB1*02:01 allele by forming hydrogen bonds with the amino acids Glu12, Leu101A, hydrophobic bonds with the amino acids Trp2, Phe4, Tyr6, Gly10, Glu12, Gln45A, Leu97A, Leu101A, Leu104A, Phe144A, Pro146A, Pro170A, Tyr174A, and pi-pi bonds with the aromatic amino acid Trp2, Phe4, Tyr6, Phe144A, Tyr174A. It interacts with the HLA-DQA1*01:02/DQB1*06:02 allele by forming hydrogen bonds with the amino acids Glu12, Asn8A, Asn95A, Leu96A, Thr109A, Asn110A, hydrophobic bonds with the amino acids Trp2, Phe4, Tyr6, Gly10, Glu12, Gly14, Gln40A, Phe41A, Ala92A, Tyr103A, Thr106A, Ala107A, Ala108A, and pi-pi bonds with the aromatic amino acid Trp2, Phe4, Tyr6, Tyr103A. It interacts with the HLA-DRB1 allele by forming hydrogen bonds with the amino acids Glu12, Val40B, Cys44B, Tyr59B, Tyr107B, Asn111B, Gln178B, hydrophobic bonds with the amino acids Trp2, Phe4, Tyr6, Gly10, Glu12, Lys41B, His42B, Tyr107B, Asn111B, Tyr112B, Trp182B, and pi-pi bonds with the aromatic amino acid Trp2, Phe4, Tyr6. The Nucleocapsid peptide (**RWYFYYLGTGPEAGL**) interacts most effectively with the HLA-DQA1*01:02/DQB1*06:02 allele, hence it showed the highest binding affinity to it (Figures 7 and 8).

Concerning the two Nucleocapsid peptides (**RWYFYYLGTGPEAGL**) and (**PRWYFYYLGTGPEAG**) in the interaction with the MHC II alleles HLA- DPA1*01:03/DPB1*02:01 and HLA-DRB1, the 2D and 3D interaction results showed that the peptide (**PRWYFYYLGTGPEAG**) more effectively with the HLA- DPA1*01:03/DPB1*02:01 allele and the peptide (**RWYFYYLGTGPEAGL**) more effectively with the HLA-DRB1 allele, however, they differ only in the first and last amino acid (Figures 9 and 10).

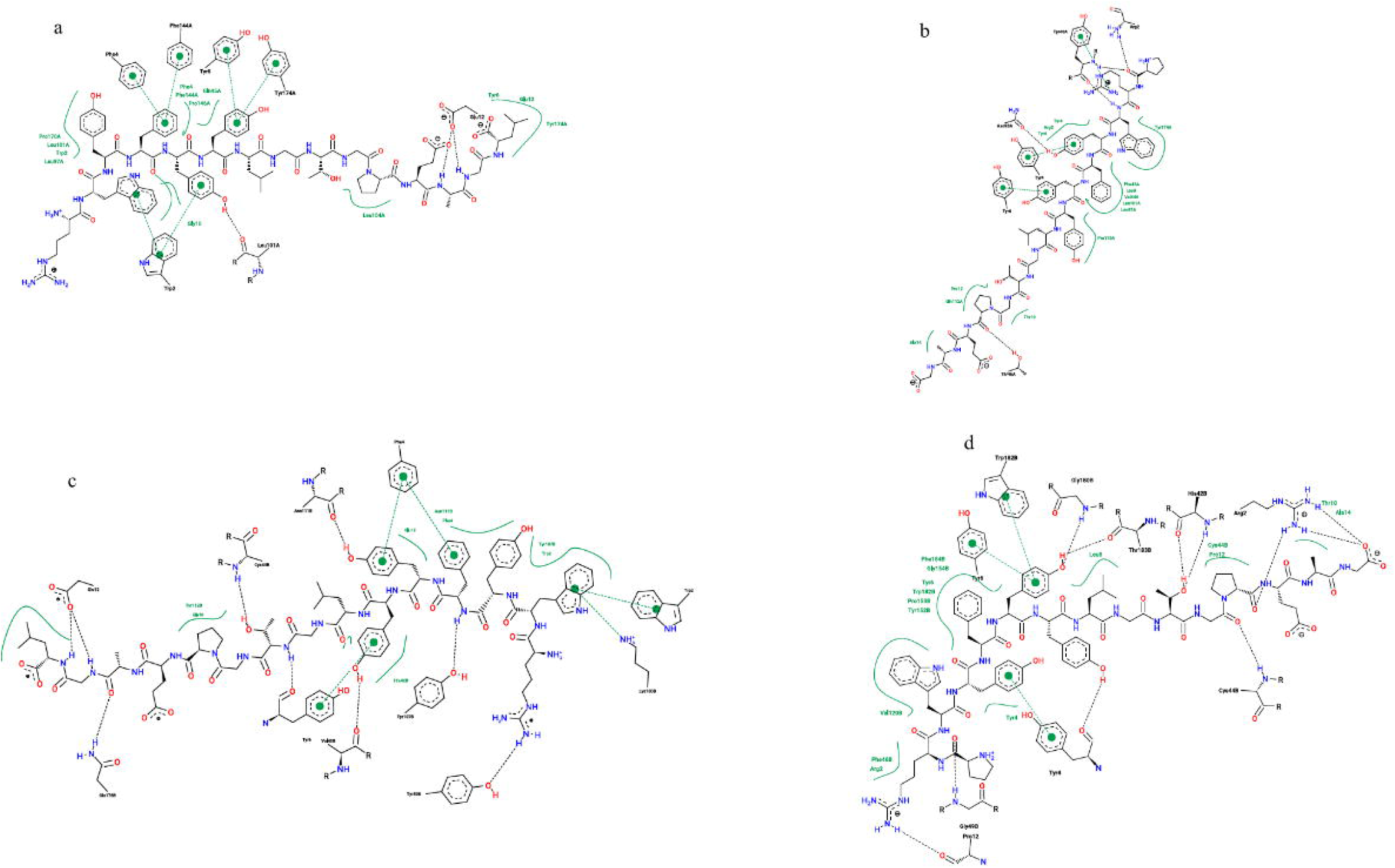

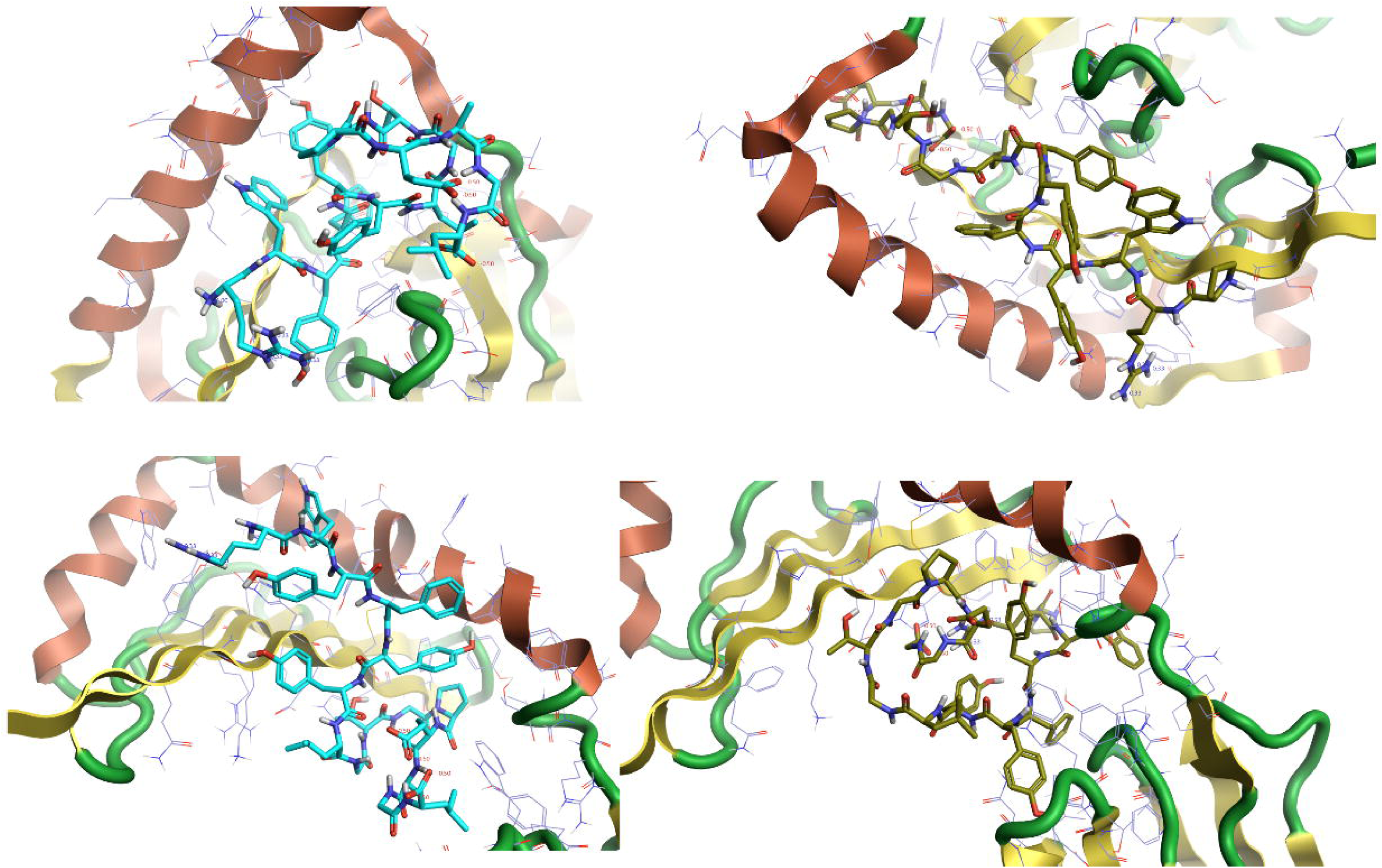

In comparison between the Spike and Nucleocapsid peptides, the Spike peptide (**FTISVTTEI**) showed a higher binding affinity to the MHC I HLA-B1503 allele. The Nucleocapsid peptides (**KTFPPTEPK**) and (**RWYFYYLGTGPEAGL**) showed a higher binding affinity to the MHC I HLA-A0202 allele and the three MHC II alleles HLA- DPA1*01:03/DPB1*02:01, HLA-DQA1*01:02/DQB1*06:02, HLA-DRB1, respectively, however, the total population coverage of the peptides **FTISVTTEI** and **KTFPPTEPK** is not high (Table 10).

**Table 10.**
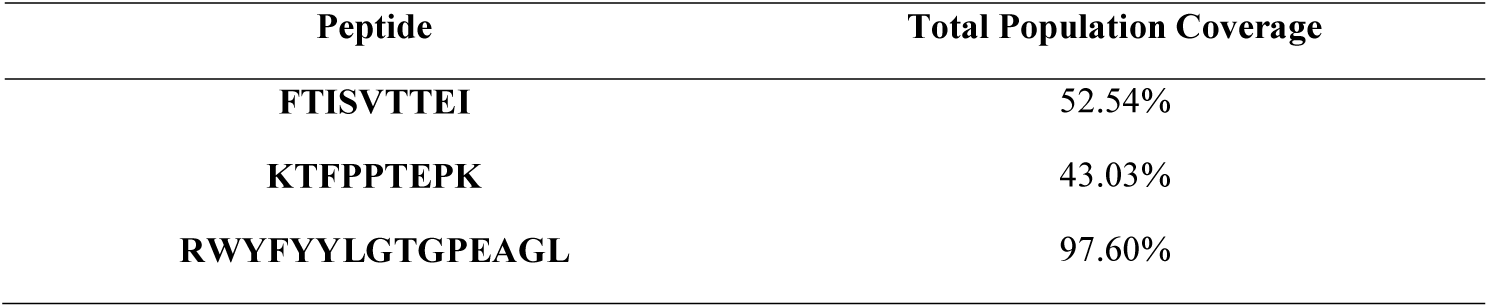
the total population coverage of the best predicted MHC peptides

Joshi A, *et al*. in their predictive COVID-19 Peptide-based vaccine study found that the ORF-7A protein’s peptide (**ITLCFTLKR**) binds most effectively with the MHC I HLA alleles HLA-A*11:01, HLA-A*68:01 [13]. Enayatkhani M, *et al*. in their predictive study included Nucleocapsid N, but they studied its interaction with the MHC I HLA-A*11:01 allele [11]. Kalita P, *et al*. included the Nucleocapsid protein and Spike glycoprotein. They used the predicted Peptides from Nucleocapsid, Spike, and Membrane glycoprotein to design a subunit vaccine [12]. Singh A, *et al*. used the Nucleocapsid protein to design multi-Peptides vaccine [14]. Since we didn’t include the two alleles HLA-A*11:01, HLA-A*68:01 in our study and didn’t design subunit or multi-Peptides vaccine, the logical comparison will not be applied.

Besides the binding with the MHC molecules, the predicted peptide must be non-toxic and non-allergen, hence, their safety was predicted using the AllergenFP v.1.0 [22], AllerCatPro [23] v. 1.7, ToxinPred [24] servers. The result showed that all the peptides were non-toxic. The AllerCatPro v. 1.7 [23] server results showed there is no evidence about the allergicity of all peptides, however, the AllergenFP v.1.0 [22] server predicts Spike peptide (**EVFNATRFASVYAWN**) as an allergen (Tables 8 and 9).

## Conclusion

A potential COVID-19 Peptide-based vaccine was predicted from the Nucleocapsid phosphoprotein (N) and Spike Glycoprotein (S) via the Immunoinformatics approach. The Spike peptide peptides **SQCVNLTTRTQLPPAYTNSFTRGVY** is predicted to have the highest binding affinity to the B-Cells. The Spike peptide **FTISVTTEI** has the highest binding affinity to the MHC I HLA-B1503 allele. The Nucleocapsid peptides **KTFPPTEPK** and **RWYFYYLGTGPEAGL** have the highest binding affinity to the MHC I HLA-A0202 allele and the three MHC II alleles HLA-DPA1*01:03/DPB1*02:01, HLA- DQA1*01:02/DQB1-*06:02, HLA-DRB1, respectively. Furthermore, those peptides were predicted as non-toxic and non-allergen. Therefore, the combination of those peptides is predicted to stimulate better immunological responses.

Since the study is an in silico predictive work, further experimental studies are recommended to validate the obtained results.

## Supporting information

Supplementary file S2 -TCELLS

## References

[1] J. Gonzalez, P. Gomez-Puertas, D. Cavanagh, A. Gorbalenya, and L. Enjuanes, “A comparative sequence analysis to revise the current taxonomy of the family Coronaviridae,” Archives of virology, vol. 148, no. 11, pp. 2207–2235, 2003.

[2] W. H. Organization, “Novel Coronavirus (2019-nCoV): situation report, 3,” 2020.

[3] K. McIntosh, “Coronavirus disease 2019 (COVID-19): Epidemiology, virology, clinical features, diagnosis, and prevention. Uptodate 2020,” ed, 2020.

[4] J. Zheng, “SARS-CoV-2: an Emerging Coronavirus that Causes a Global Threat,” (in eng), Int J Biol Sci, vol. 16, no. 10, pp. 1678–1685, 2020, DOI: 10.7150/ijbs.45053.

[5] E. C. f. D. P. a. Control. “COVID-19 situation update worldwide, as of 5 May 2020.” https://www.ecdc.europa.eu/en/geographical-distribution-2019-ncov-cases (accessed 6/5/2020, 2020).

[6] T. W. H. O. (WHO). “WHO Solidarity Trial – Accelerating a safe and effective COVID-19 vaccine.” https://www.who.int/emergencies/diseases/novel-coronavirus-2019/global-research-on-novel-coronavirus-2019-ncov/solidarity-trial-accelerating-a-safe-and-effective-covid-19-vaccine (accessed 6/5/2020, 2020).

[7] T. T. Le et al., “The COVID-19 vaccine development landscape,” Nat Rev Drug Discov, 2020.

[8] J. F.-W. Chan et al., “Genomic characterization of the 2019 novel human-pathogenic coronavirus isolated from a patient with atypical pneumonia after visiting Wuhan,” Emerging microbes & infections, vol. 9, no. 1, pp. 221–236, 2020.

[9] W. Li, M. D. Joshi, S. Singhania, K. H. Ramsey, and A. K. Murthy, “Peptide Vaccine: Progress and Challenges,” Vaccines, vol. 2, no. 3, pp. 515–536, 2014, DOI: 10.3390/vaccines2030515.

[10] E. Petitdidier et al., “Peptide-based vaccine successfully induces protective immunity against canine visceral leishmaniasis,” npj Vaccines, vol. 4, no. 1, p. 49, 2019/11/29 2019, DOI: 10.1038/s41541-019-0144-2.

[11] M. Enayatkhani et al., “Reverse vaccinology approach to design a novel multi-epitope vaccine candidate against COVID-19: an in silico study,” Journal of Biomolecular Structure and Dynamics, pp. 1-16, 2020, DOI: 10.1080/07391102.2020.1756411.

[12] P. Kalita, A. K. Padhi, K. Zhang, and T. Tripathi, “Design of a Peptide-Based Subunit Vaccine against Novel Coronavirus SARS-CoV-2,” ed: Preprints, 2020.

[13] A. Joshi, B. C. Joshi, M. A.-u. Mannan, and V. Kaushik, “Epitope based vaccine prediction for SARS-COV-2 by deploying immuno-informatics approach,” Informatics in Medicine Unlocked, vol. 19, p. 100338, 2020/01/01/2020, DOI: https://doi.org/10.1016/j.imu.2020.100338.

[14] A. Singh, M. Thakur, L. K. Sharma, and K. Chandra, “Designing a multi-epitope peptide-based vaccine against SARS-CoV-2,” bioRxiv, p. 2020.04.15.040618, 2020, DOI: 10.1101/2020.04.15.040618.

[15] N. R. Coordinators, “Database resources of the national center for biotechnology information,” Nucleic acids research, vol. 45, no. Database issue, p. D12, 2017.

[16] J. Robinson, D. J. Barker, X. Georgiou, M. A. Cooper, P. Flicek, and S. G. E. Marsh, “IPD-IMGT/HLA Database,” Nucleic Acids Research, vol. 48, no. D1, pp. D948–D955, 2019, DOI: 10.1093/nar/gkz950.

[17] D. G. H. Julie D. Thompson, and Toby J. Gibson, “CLUSTAL W: improving the sensitivity of progressive multiple sequence alignment through sequence weighting, position-specific gap penalties, and weight matrix choice,” Nucleic Acids Research, vol. 22, no. 22, pp. 4673–80, 1994.

[18] T. A. Hall, “BioEdit: a user-friendly biological sequence alignment editor and analysis program for Windows 95/98/NT,” Nucleic Acids Symposium Series, vol. 41, pp. 95–98, 1999, DOI: CiteULike-article-id:691774.

[19] R. Vita et al., “The immune epitope database (IEDB) 3.0,” Nucleic acids research, vol. 43, no. Database issue, pp. D405–D412, 2015, DOI: 10.1093/nar/gku938.

[20] J. E. P. Larsen, O. Lund, and M. Nielsen, “Improved method for predicting linear B-cell epitopes,” Immunome Research, vol. 2, no. 1, p. 2, 2006.

[21] E. A. Emini, J. V. Hughes, D. Perlow, and J. Boger, “Induction of hepatitis A virus-neutralizing antibody by a virus-specific synthetic peptide,” Journal of virology, vol. 55, no. 3, pp. 836–839, 1985.

[22] L. N. Ivan Dimitrov, Irini Doytchinova, and Ivan Bangov, “AllergenFP: allergenicity prediction y descriptor fingerprints,” Bioinformatics, vol. 30, no. 6, pp. 846–851, 2013.

[23] S. Maurer-Stroh et al., “AllerCatPro—prediction of protein allergenicity potential from the protein sequence,” Bioinformatics, vol. 35, no. 17, pp. 3020–3027, 2019, DOI: 10.1093/bioinformatics/btz029.

[24] S. Gupta et al., “In silico approach for predicting toxicity of peptides and proteins,” PloS one, vol. 8, no. 9, pp. e73957–e73957, 2013, DOI: 10.1371/journal.pone.0073957.

[25] J. K. Torsten Schwede, Nicolas Guex, and Manuel C. Peitsch, “SWISS-MODEL: an automated protein homology-modeling server,” Nucleic Acids Research, vol. 31, no. 13, pp. 3381–3385, 2003.

[26] K. e. al., “The Phyre2 web portal for protein modeling, prediction, and analysis,” Nature Protocols, vol. 10, no. 845-858, 2015.

[27] G. T. Pettersen EF, Huang CC, Couch GS, Greenblatt DM, Meng EC, Ferrin TE, “UCSF Chimera--a visualization system for exploratory research and analysis,” J Comput Chem., vol. 25, no. 13, pp. 1605–1612, 2004.

[28] v. Flare, Cresset®, Litlington, Cambridgeshire, UK, http://www.cresset-group.com/flare/;

[29] Cheeseright, T.; Mackey, M.; Rose, S.; Vinter, “A. Molecular Field Extrema as Descriptors of Biological Activity:LJ Definition and Validation.,” J. Chem. Inf. Model., vol. 46, no. 2, pp. 665–676., 2006.

[30] X. Xu, C. Yan, and X. Zou, “MDockPeP: An ab-initio protein–peptide docking server,” Journal of Computational Chemistry, vol. 39, no. 28, pp. 2409–2413, 2018, DOI: 10.1002/jcc.25555.

[31] P. Zhou, B. Jin, H. Li, and S.-Y. Huang, “HPEPDOCK: a web server for blind peptide-protein docking based on a hierarchical algorithm,” (in eng), Nucleic acids research, vol. 46, no. W1, pp. W443–W450, 2018, DOI: 10.1093/nar/gky357.

[32] M. Remmert, A. Biegert, A. Hauser, and J. Söding, “HHblits: lightning-fast iterative protein sequence searching by HMM-HMM alignment,” Nature Methods, vol. 9, no. 2, pp. 173–175, 2012/02/01 2012, DOI: 10.1038/nmeth.1818.

[33] W. R. Pearson and D. J. Lipman, “Improved tools for biological sequence comparison,” (in eng), Proceedings of the National Academy of Sciences of the United States of America, vol. 85, no. 8, pp. 2444–2448, 1988, DOI: 10.1073/pnas.85.8.2444.

[34] M. A. Martí-Renom, A. C. Stuart, A. Fiser, R. Sánchez, F. M. and, and A. Šali, “Comparative Protein Structure Modeling of Genes and Genomes,” Annual Review of Biophysics and Biomolecular Structure, vol. 29, no. 1, pp. 291–325, 2000, DOI: 10.1146/annurev.biophys.29.1.291.

[35] H. M. Berman et al., “The Protein Data Bank,” (in eng), Nucleic acids research, vol. 28, no. 1, pp. 235–242, 2000, DOI: 10.1093/nar/28.1.235.

[36] K. S. a. M. Rarey, “PoseView--molecular interaction patterns at galance,” Journal of Cheminformatic., vol. 2, no. 1, p. 50, 2010.

[37] R. Fährrolfes et al., “ProteinsPlus: a web portal for structure analysis of macromolecules,” Nucleic Acids Research, vol. 45, no. W1, pp. W337–W343, 2017, DOI: 10.1093/nar/gkx333 %J Nucleic Acids Research.

[38] N. Lurie, M. Saville, R. Hatchett, and J. Halton, “Developing Covid-19 Vaccines at Pandemic Speed,” New England Journal of Medicine, 2020, DOI: 10.1056/NEJMp2005630.

[39] C. Rueckert and C. A. Guzmán, “Vaccines: From Empirical Development to Rational Design,” PLOS Pathogens, vol. 8, no. 11, p. e1003001, 2012, DOI: 10.1371/journal.ppat.1003001.

[40] U. K. Adhikari, M. Tayebi, and M. M. Rahman, “Immunoinformatics Approach for Epitope-Based Peptide Vaccine Design and Active Site Prediction against Polyprotein of Emerging Oropouche Virus,” Journal of Immunology Research, vol. 2018, p. 6718083, 2018/10/08 2018, DOI: 10.1155/2018/6718083.

[41] I. A. Resource. “Antibody Epitope Prediction – Tutorial.” http://tools.iedb.org/bcell/help/ (accessed 5/9/2020, 2020).

[42] “Chapter 3 – Immunogenicity and Antigenicity,” in Immunology for Pharmacy, D. K. Flaherty Ed. Saint Louis: Mosby, 2012, pp. 23–30.

[43] K. L. Rock, E. Reits, and J. Neefjes, “Present Yourself! By MHC Class I and MHC Class II Molecules,” (in eng), Trends Immunol, vol. 37, no. 11, pp. 724–737, 2016, DOI: 10.1016/j.it.2016.08.010.

[44] Z. Cournia, B. Allen, and W. Sherman, “Relative binding free energy calculations in drug discovery: recent advances and practical considerations,” Journal of chemical information and modeling, vol. 57, no. 12, pp. 2911–2937, 2017.

[45] A. Nguyen et al., “Human leukocyte antigen susceptibility map for SARS-CoV-2,” medRxiv, p. 2020.03.22.20040600, 2020, DOI: 10.1101/2020.03.22.20040600.

[46] M. Wieczorek et al., “Major Histocompatibility Complex (MHC) Class I and MHC Class II Proteins: Conformational Plasticity in Antigen Presentation,” (in English), Frontiers in Immunology, Review vol. 8, no. 292, 2017-March-17 2017, DOI: 10.3389/fimmu.2017.00292.

[47] T. M. Holling, E. Schooten, and P. J. van Den Elsen, “Function and regulation of MHC class II molecules in T-lymphocytes: of mice and men,” Human Immunology, vol. 65, no. 4, pp. 282–290, 2004.

