## Supplementary material for "A Multiple Peptides Vaccine against nCOVID-19 Designed from the Nucleocapsid phosphoprotein (N) and Spike Glycoprotein (S) via the Immunoinformatics Approach": Supplmentary file S1

Supplementary Table S1 the Nucleocapside phosphoprotein sequences retrieved from the NCBI database [1].

| **Accession number** | **Country** | **Collection Date** |
| --- | --- | --- |
| YP_009724390.1 | China | DEC-2019 |
| QHR84449.1 | Australia | 25-JAN-2020 |
| QHN73810.1 | China | 11-JAN-2020 |
| QHN73795.1 | China | 10-JAN-2020 |
| QHO60594.1 | USA | 19-JAN-2020 |
| QHQ82464.1 | USA | 22-JAN-2020 |
| QHQ71973.1 | USA | 22-JAN-2020 |
| QHQ71963.1 | USA | 23-JAN-2020 |
| QHO62112.1 | China | 02-JAN-2020 |
| QHO62107.1 | China | 02-Jan-2020 |

Supplementary Table S2 the Spike glycoprotein sequences retrieved from the NCBI database [1].

| **Accession number** | **Country** | **Collection Date** |
| --- | --- | --- |
| QIM47483.1 | Spain | March 2020 |
| QIM47464.1 | Spain | March 2020 |
| QHD43423.2 | China | 2020 |
| QIK50455.1 | Viet Nam | MAR-2020 |
| QIK50445.1 | Viet Nam | MAR-2020 |
| QIK50435.1 | USA | MAR-2020 |
| QHU79181.1 | Finland | FEB-2020 |
| QIK50424.1 | Taoyuan | MAR-2020 |
| QIK02971.1 | USA | MAR-2020 |
| QIK02961.1 | USA | MAR-2020 |
| QIK02951.1 | USA | MAR-2020 |
| QIJ96530.1 | USA | MAR-2020 |
| QIJ96520.1 | USA | MAR-2020 |
| QIJ96510.1 | USA | MAR-2020 |
| QIJ96500.1 | USA | MAR-2020 |
| QIJ96490.1 | USA | MAR-2020 |
| QIJ96480.1 | USA | MAR-2020 |
| QIJ96470.1 | USA | MAR-2020 |
| QIE07488.1 | China | Jan-2020 |
| QIE07478.1 | China | Jan-2020 |
| QIE07468.1 | China | Jan-2020 |
| YP_009724397.2 | China | Dec-2019 |
| BCB15098.1 | Japan | Jan-2020 |
| QII87848.1 | USA | Mar-2020 |
| QII87838.1 | USA | Mar-2020 |
| QII87826.1 | USA | Mar-2020 |
| QII87814.1 | USA | Feb-2020 |
| QII87802.1 | USA | Feb-2020 |
| QII87790.1 | USA | Feb-2020 |
| QII57345.1 | USA | Feb-2020 |
| QII57335.1 | USA | Feb-2020 |
| QII57325.1 | USA | Feb-2020 |
| QII57315.1 | USA | Feb-2020 |
| QII57305.1 | USA | Feb-2020 |
| QII57295.1 | USA | Feb-2020 |
| QII57285.1 | USA | Feb-2020 |
| QII57275.1 | USA | Feb-2020 |
| QII57265.1 | USA | Feb-2020 |
| QII57255.1 | USA | Feb-2020 |
| QII57245.1 | USA | Feb-2020 |
| QII57235.1 | USA | Feb-2020 |
| QII57225.1 | USA | Feb-2020 |
| QII57215.1 | USA | Feb-2020 |
| QII57205.1 | USA | Feb-2020 |
| QII57195.1 | USA | Feb-2020 |
| QII57185.1 | USA | Feb-2020 |
| QII57175.1 | USA | Feb-2020 |
| QIA98561.1 | Italy | Jan-2020 |
| QIH55228.1 | USA | Feb-2020 |
| QIA98590.1 | India | Jan-2020 |
| QHS34553.1 | India | Jan-2020 |
| QIG56001.1 | Brazil | Feb-2020 |
| QIG55986.1 | Viet Nam | Feb-2020 |
| BCA87378.1 | Japan | Feb-2020 |
| BCA87368.1 | Japan | Feb-2020 |
| QIE07458.1 | China | Feb-2020 |
| QID98801.1 | USA | Feb-2020 |
| QID21075.1 | USA | Feb-2020 |
| QID21065.1 | USA | Feb-2020 |
| QID21055.1 | USA | Feb-2020 |
| QIC53211.1 | Sweden | Feb-2020 |
| QIC53211.1 | Sweden | Feb-2020 |
| QIC50516.1 | China | Feb-2020 |
| QIC50515.1 | China | Jan-2020 |
| QIC50514.1 | China | Jan-2020 |
| QIC50513.1 | China | Jan-2020 |
| QIC50512.1 | China | Jan-2020 |
| QIC50511.1 | China | Jan-2020 |
| QIC50510.1 | China | Jan-2020 |
| QIC50509.1 | China | Jan-2020 |
| QIC50508.1 | China | Jan-2020 |
| QIC50507.1 | China | Jan-2020 |
| QIB84680.1 | Nepal | Jan-2020 |
| QIA98613.1 | Taiwan | Feb-2020 |
| QIA98602.1 | Taiwan | Jan-2020 |
| QIA20052.1 | China | Jan-2020 |
| QHZ87599.1 | USA | Jan-2020 |
| QHZ87589.1 | USA | Jan-2020 |
| QHZ00406.1 | USA | Jan-2020 |
| QHZ00396.1 | USA | Jan-2020 |
| QHZ00386.1 | South Korea | Jan-2020 |
| QHW06066.1 | USA | Jan-2020 |
| QHW06056.1 | USA | Jan-2020 |
| QHW06046.1 | USA | Jan-2020 |
| QHU79211.1 | USA | Jan-2020 |
| QHU79201.1 | USA | Jan-2020 |
| QHU36871.1 | China | Jan-2020 |
| QHU36861.1 | China | Dec-2019 |
| QHU36851.1 | China | Dec-2019 |
| QHU36841.1 | China | Dec-2019 |
| QHU36831.1 | China | Dec-2019 |
| QHR84456.1 | Australia | Jan-2020 |
| QHQ82471.1 | USA | Jan-2020 |
| QHQ71980.1 | USA | Jan-2020 |
| QHQ71970.1 | USA | Jan-2020 |
| QHO62884.1 | USA | Jan-2020 |
| QHO60601.1 | USA | Jan-2020 |
| QHN73817.1 | China | Jan-2020 |
| QHN73802.1 | China | Jan-2020 |

### **References**

[1]	N. R. Coordinators, "Database resources of the national center for biotechnology information," *Nucleic acids research,* vol. 45, no. Database issue, p. D12, 2017.
